# Single-cell analysis identifies TCF4 and ID3 as a molecular switch of mammary epithelial stem cell differentiation

**DOI:** 10.1101/2020.08.16.249854

**Authors:** Koon-Kiu Yan, Erin Nekritz, Bensheng Ju, Xinran Dong, Rachel Werner, Dayanira Alsina-Beauchamp, Celeste Rosencrance, Partha Mukhopadhyay, Qingfei Pan, Andrej Gorbatenko, Liang Ding, Yanyan Wang, Chenxi Qian, Hao Shi, Bridget Shaner, Sivaraman Natarajan, Hongbo Chi, John Easton, Jose Silva, Jiyang Yu

## Abstract

It is well known that the expansion of the mammary epithelium during the ovarian cycles in female mammals is supported by the transient increase in mammary epithelial stem cells (MaSCs). However, dissecting the molecular mechanisms that govern MaSC function and differentiation is poorly understood due to the lack of standardized methods for their identification and isolation.

The development of robust single-cell mRNA sequencing () technologies and the computational methods to analyze them provides us with novel tools to approach the challenge of studying MaSCs in a completely unbiased way without. Here, we have performed the largest scRNA-seq analysis of individual mammary epithelial cells (~70,000 cells). Our study identified a distinct cell population presenting molecular features of MaSCs.

Importantly, further purification and additional in-depth single-cell analysis of these cells revealed that they are not a fully homogenous entity. Instead, we identified three subpopulations representing early stages of lineage commitment. By tracking their molecular evolution through single-cell network analysis we found that one of these subpopulations represents bipotent MaSCs from which luminal and basal lineages diverge. Importantly, we also confirmed the presence of these cells in human mammary glands. Finally, through expression and network analysis studies, we have uncovered transcription factors that are activated early during lineage commitment. These data identified E2-2 (Tcf4) and ID3 as a potential molecular switch of mammary epithelial stem cell differentiation.

## INTRODUCTION

The mammary gland is a dynamic organ that undergoes extensive tissue remodeling over the female lifespan^1–4^. In the adult female, expansion and regression of the basal and luminal bilayers occur regularly during the ovarian cycles. Additionally, massive epithelial expansion and generation of specialized secretory cells occur during pregnancy, resulting in the generation of specialized secretory glands. In all of these stages, epithelial expansion is supported by a transient increase in the mammary epithelial stem cell (MaSC) population^5–7^.

Although the existence of MaSCs is well known^8,9^, two main limitations have handicapped the dissection of the molecular mechanisms that govern their function and differentiation. The first issue is the limited ability to identify and isolate MaSCs. For this, fluorescence-activated cell sorting (FACS) using antibodies against membrane markers followed by transplantation into cleared mammary fat pads has been used to examine MaSC potential. However, these assays represent non-physiological conditions where lineage-committed cells demonstrate facultative stem cell activity that does not occur *in vivo^8,10^*. Thus, during recent years, the study of MaSCs has turned to lineage tracing experiments. Here, cells of interest are labeled *in vivo* at an initial time point, and the progeny derived from these cells is tracked over time ^5,11–13^. Unfortunately, such experiments have generated conflicting results. Although some studies have reported the existence of luminal and basal unipotent MaSCs^14^, others have found bipotent MaSCs that replenish both lineages^5,11–13^. Importantly, both FACS and lineage tracing experiments rely on the use of predetermined markers or promoters and, consequently, are subject to bias.

The second limitation in studying MaSCs involves the methods used to identify genes or pathways involved in lineage commitment. Generally, molecular features (such as mRNA profiling) of a MaSC-enriched population are compared with those of differentiated luminal and basal cells^12,14–16^. Importantly, however, this type of comparison is handicapped by the very nature of fate specification. Molecularly, differentiation is a continuous process in which a few signals start a cascade that leads to profound changes observed between uncommitted cells and the fully differentiated progeny. However, the initiating and subsequent signals do not necessarily accumulate, and instead can appear and disappear without being detected at the time when cells are being processed. Thus, delineating the molecular hierarchy that regulates lineage commitment of the mammary epithelium requires the identification of pure MaSC populations and elucidation of the molecular changes that occur early during the process.

The rapid development of robust single-cell mRNA sequencing (scRNA-seq) technologies and the computational methods to analyze them provides us with novel tools to approach the challenge of studying MaSCs in a completely unbiased way without any selection of predetermined markers or perturbations. Thus, by coupling scRNA-seq with state-of-the-art systems biology^17,18^ we have analyzed over 70,000 individual mammary epithelial cells. Our study clearly identified a distinct population of cells characterized by the expression of the protein C receptor, which has recently been shown to be expressed in multipotent cells in several tissues^13,19,20^. Importantly, additional in-depth single-cell analysis of these cells identified three subpopulations representing an early stage of lineage commitment. By tracking their molecular evolution through single-cell network analysis we found that one of these subpopulations represents bipotent MaSCs from which luminal and basal linages diverge. Importantly, we also confirmed the presence of these cells in human mammary glands after performing similar studies. Finally, through expression and network analysis studies, we have uncovered transcription factors that are activated early during lineage commitment. These data identified E2-2 (*Tcf4*) and ID3 as a molecular switch of mammary epithelial stem cell differentiation.

## RESULTS

### Single-cell studies of the mouse mammary gland identify distinct cell subpopulations

ScRNA‐seq is a powerful method to identify cell subtypes and track the trajectories of cell lineages^21,22^. We performed scRNA-seq on mammary epithelial cells (MECs) isolated at different stages of the female mouse development: puberty (6 weeks), virgin adult (4 months) and old (24 months) (Fig. 1A). Because of the well-known impact of the ovarian hormones during the estrous cycle^4,6,7^, we further separated adult females into estrus and diestrus groups. For isolation, non-epithelial cells were removed by antibodies against CD31^−^, CD45^−^, Ter119^−^ and BP1^−^ ^23–25^ and EpCAM^+^epithelial cells were isolated by FACS. In total, we evaluated ~20,000 MECs per time point (>70,000 cells total) using a droplet‐based platform (10X Genomics)^21,26^, which is one of the largest scRNA-seq studies ever performed. Clustering analysis of all sequenced cells based on t-distributed stochastic neighbor embedding (tSNE) plotting^21,22^ identified three of the major clusters as basal, luminal estrogen receptor-positive (ER+) and luminal estrogen receptor-negative (ER-) cells (Fig. 1A–1C). Remarkably, an additional small population also emerged (Fig.1A). This population did not present high levels of basal or luminal markers but instead showed the distinct expression of the protein C receptor (PROCR) (Fig. 1B–C) which has been linked to multipotent stem cells in endothelium^19^, keratinocytes^20^, and mammary epithelial cells^13^. In agreement with previous reports^13^, we found that PROCR+ cells express high levels of mesenchymal genes such as Vim, Cdh5, Zeb1, Zeb2, Twist1, and low levels of epithelial genes such as Epcam, Cdh1, Krt14, Krt5, Krt8, and Krt18. The size of the PROCR+ population ranged from 0.3 to 3% of the total mammary epithelium, and it was more prevalent during puberty (Fig. 1C and data not shown).

**Fig. 1.**
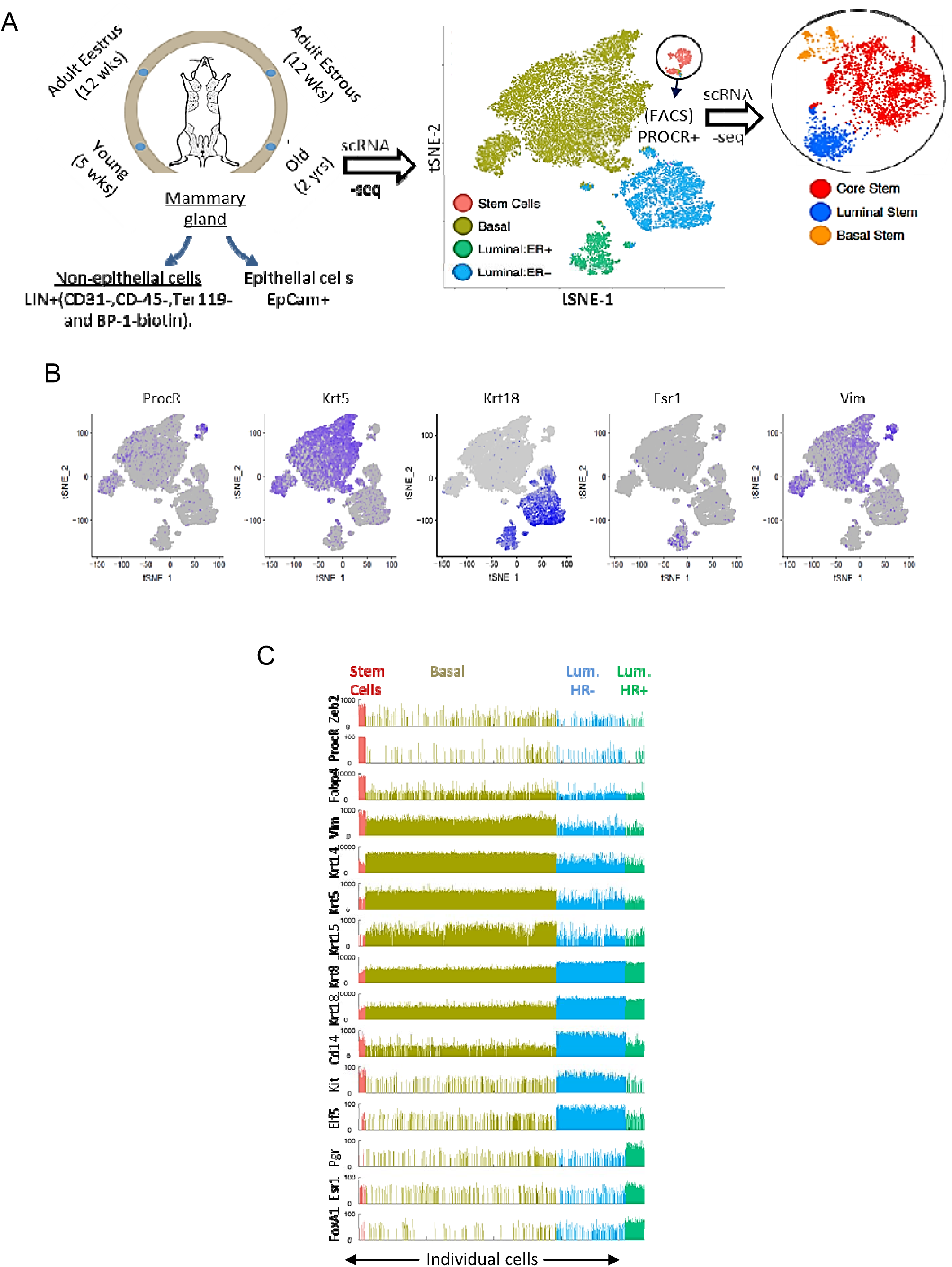
Single-cell transcriptomics of mammary epithelium: A) Purification and scRNA-seq profile of Balb/cj virgin females as described in the text. The tSNE plot shows all the different time points merged and an enlarged plot of the MaSC subpopulations. B) tSNE plot from A showing the expression of specific genes (purple). C) The bar plot shows the expression of lineage-specific markers in mammary epithelial subpopulations. Cell subpopulations are color-coded.

### PROCR+ cells consist of three subpopulations representing MaSC differentiation routes

To investigate the PROCR+ population at a very high resolution we isolated the PROCR+ population by FACS as previously described^13^ and performed scRNA-seq on ~2,000 PROCR+ cells from each time point (Fig. 1A). Remarkably we found that PROCR+ cells are not a single entity but rather consist of three distinct cell clusters (Fig. 1A) that are seen in all developmental stages (Fig.2A). To investigate the molecular characteristics of these clusters, we first defined the basal and luminal signatures by performing differential expression analysis of the scRNA-seq profiles of differentiated basal and luminal subpopulations. Then, we used the NetBID algorithm^18,27^ to infer the activities for basal or luminal signatures in each cell across all populations. We found that one of the three PROCR+ clusters presented higher luminal signature activity while another presented higher basal activity, whereas the last was undefined (Fig. 2B). For simplicity, we have designated these MaSC subtypes luminal-MaSCs, basal-MaSCs, and core-MaSCs, respectively.

**Fig. 2.**
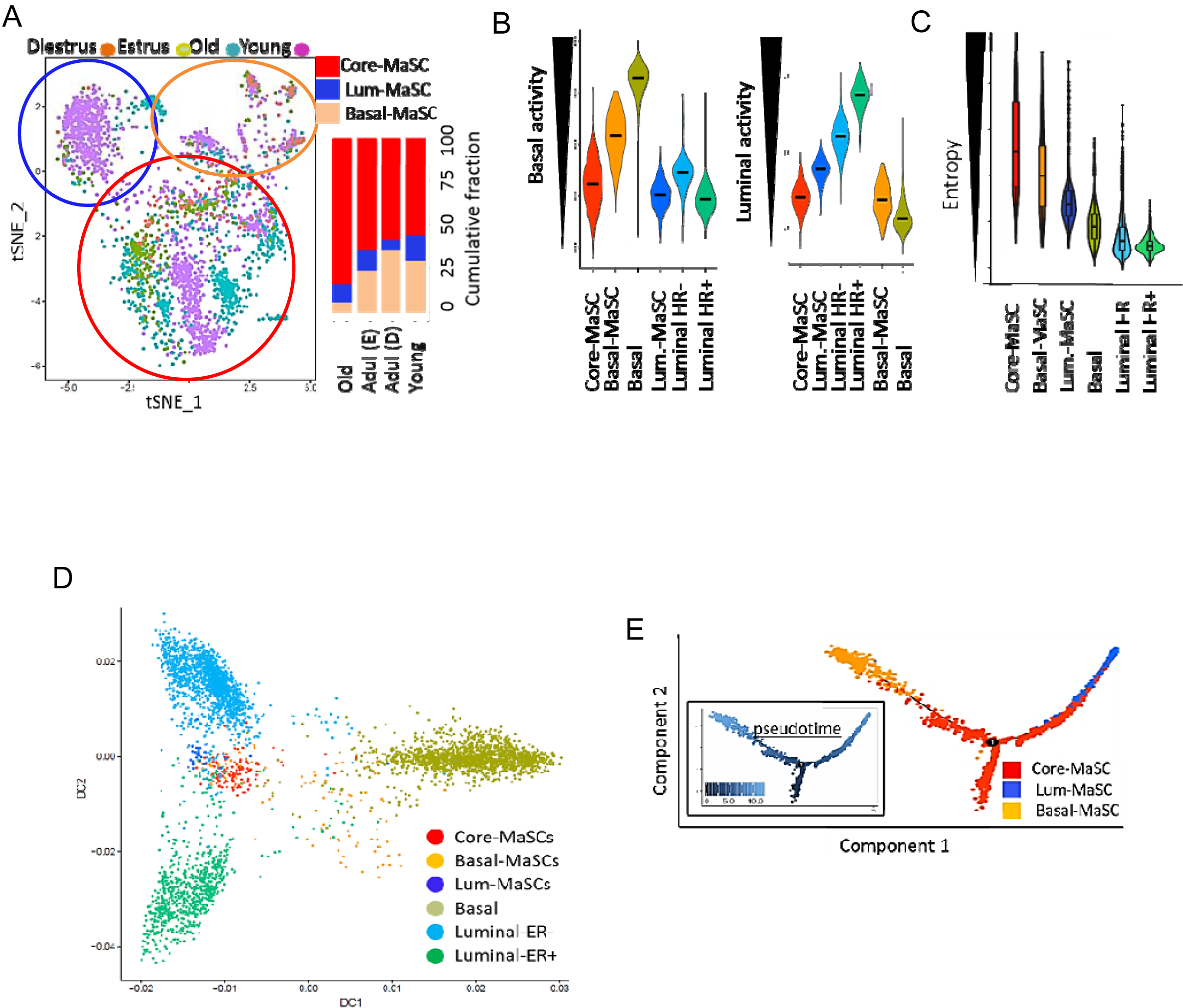
Molecular characterization of MaSC populations: A) The tSNE plot and the bar plots show the distribution of MaSCs populations in all developmental stages evaluated. B) Basal and luminal activity of MaSC subpopulations and differentiated cells and C) entropy (transcriptional plasticity) calculated as described in the text. D) Diffusion plot showing the lineage relationship of the six subpopulations. E) Pseudotime analysis showing a divergent trajectory from core-MaSCs to basal-MaSCs and luminal-MaSCs. In A and D the colored circles indicate the position of the MaSC subpopulations.

It has been reported that the differentiation potential of a cell can be approximated by computing the signaling promiscuity, or entropy, of the cell’s transcriptome, thus providing a means to identify stem cell phenotypes^28^. Interestingly, the core-MaSC subpopulation presented the highest entropy, indicative of more plasticity and therefore the highest differentiation potential (Fig. 2C). Next, we investigated the lineage relationship of these cells by generating a diffusion plot (Fig. 2D). The differentiated populations relocated to well-defined groups forming a triangle-shaped pattern. In contrast, the core-MaSCs were positioned at the geometrical center of the triangle while the basal and luminal-MaSCs extended from the core-MaSCs toward the corresponding lineage-committed cells. Finally, we dissected the differentiation trajectories of MaSCs by pseudotime analysis^29,30^. This computational method embeds scRNA-seq profiles in a low-dimensional space where the distance between adjacent cells represents the progression through a continuous but stochastic differentiation process. This analysis showed a gradual bifurcation from core-MaSCs into two branches: basal-MaSCs and luminal-MaSCs (Fig. 2E). Overall, our studies position core-MaSCs at the very top of the MaSC hierarchy.

### Single-cell studies of the human breast identify PROCR+ cells analogous to mouse MaSCs

Next, we obtained normal healthy tissue from mastectomies and performed scRNA-seq on the dissociated human MECs. Remarkably, we observed the presence of a small subpopulation highly similar to the mouse PROCR+ MaSCs (Fig. 3A). To obtain enough PROCR+ cells for further analysis, we used a FACS strategy similar to the one used in mice (Fig. 3B) and performed scRNA-seq. Here, we found that human PROCR+ cells contained three subpopulations that are analogous to the mouse core-, luminal and basal-MaSCs (Fig. 3C).

**Fig. 3.**
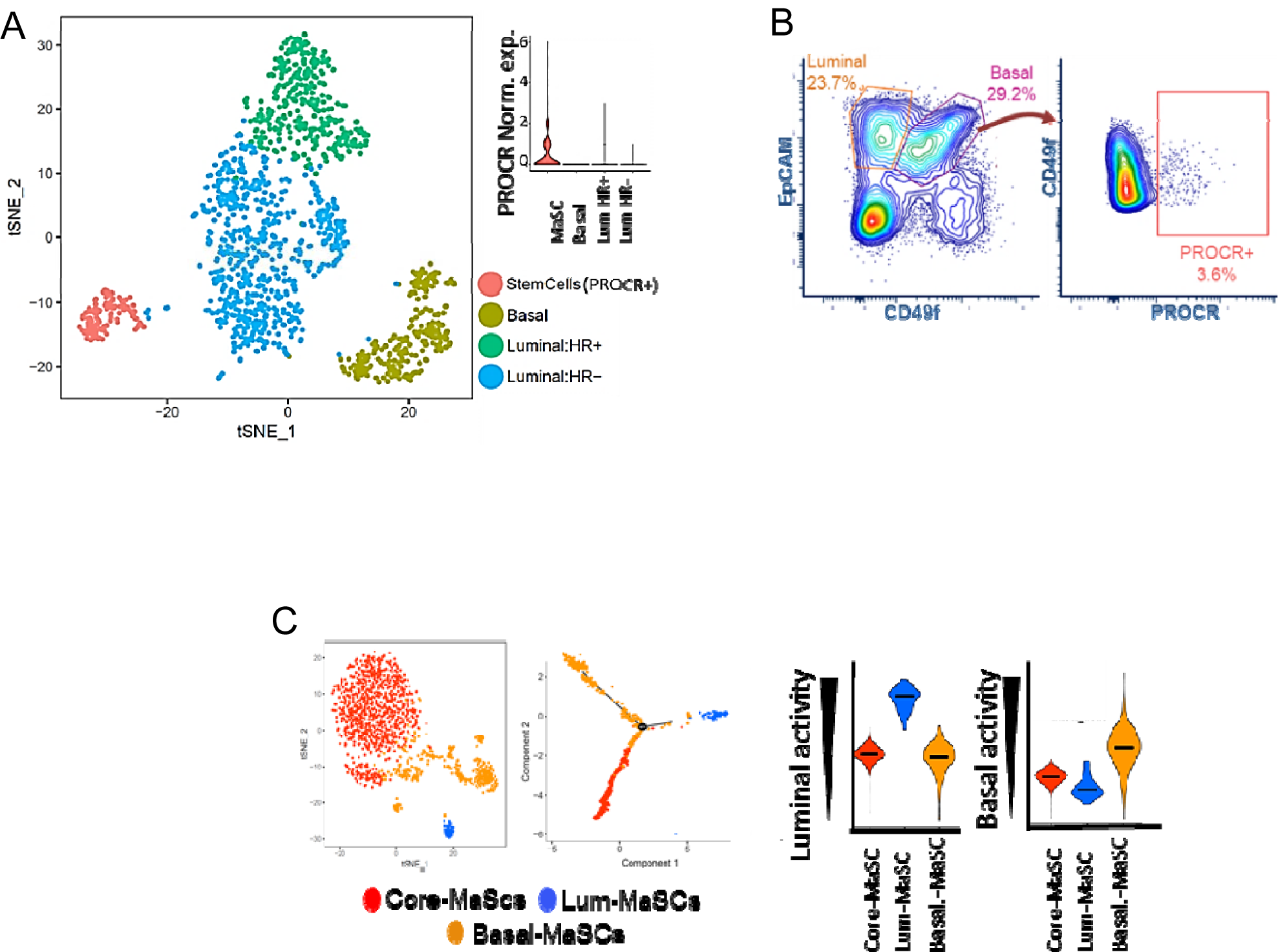
Purification of PROCR+ human cells: A) tSNE plots showing the human mammary epithelial subpopulations. The violin plot shows the expression of PROCR in each subpopulation. B) FACS profiles showing the purification of PROCR+ cells. C) tSNE-plots showing the subpopulations as well as a diffusion map and the basal and luminal activity of the purified PROCR+ MaSCs from B.

### Molecular dissection of the differentiation routes of bipotent MaSCs

ScRNA-seq analysis provides expression data for ~2,000 to 3,000 of all the expressed genes^31,32^. Although this is enough to sort cell heterogeneity, complete transcriptomic profiles are necessary for studying cell regulation. Thus, we devised a FACS strategy to isolate the core-MaSC, basal-MaSC, and luminal-MaSC subpopulations and subjected them to bulk RNA-seq (Fig. 4). To this end, we searched our scRNA-seq data to select membrane proteins specifically expressed in each of the MaSC subpopulations. We found that the core- and basal-MaSCs express high levels of CD36 and podoplanin (Pdpn) respectively, while the luminal-MaSCs do not express either of these genes (Fig. 4A). Thus, we purified PROCR+ (EpCAM^+^/CD49f^high^/PROCR^+^) cells and used used these markers to sort the MaSC subpopulations (CD36^+^ to obtain core-MaSCs, Pdpn^+^ for basal-MaSCs and CD36^−^/Pdpd^−^ for luminal-MaSCs) (Fig. 4B). As expected, PCA analysis of the sc- and the bulk RNA profiles were highly correlated, validating our FACS strategy (Fig. 4C).

**Fig. 4.**
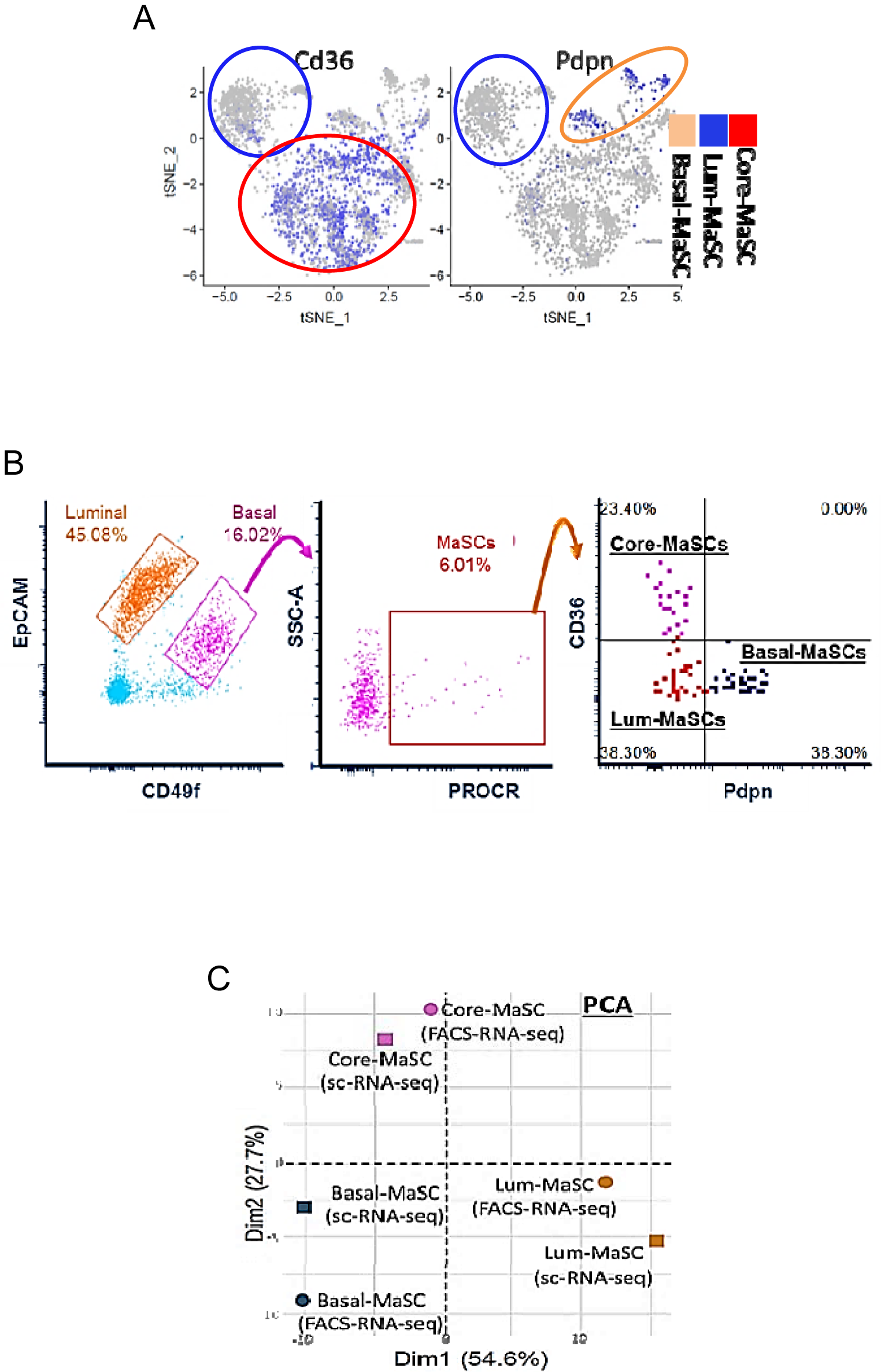
Purification of MaSC subpopulations: A) tSNE plots showing the expression of *Cd36* and *Pdpn* in MaSCs subpopulations. B) FACS plots showing the sorting strategy for isolation of MaSCs subpopulations. C) PCA plots showing the correlation between scRNA-seq and bulk RNA-seq data.

To identify key regulators of the differentiation process we used a powerful method that analyzes expression profiling called inference of transcriptional regulatory networks (TRNs)^26,33–37^. Briefly, a TRN consists of a web of connections (network) between transcription factors (TFs) and their targets. A TF-gene connection can show a positive (activator) or a negative (repressor) correlation. The activity of a TF is calculated based on the expression of its targets and is represented numerically. Subsequently, the TRN is inferred by integrating all the TF-genes hubs. To delineate TRNs, we used the SJARACNe^17^ algorithm (an improved implementation of ARACNe^38^ that has been used extensively in normal and cancer cells^26,33–37^). Then, we used the MaSC-specific TRN to identify the Master Regulators (MRs) of each MaSC subpopulation. A MR is defined as a TF that is differentially active among the populations studied^35–37,39–45^. For this, we used NetBID^27^ to calculate the activity scores of each TF in each MaSC subpopulation. To quantify the significance of each activity score we performed gene set enrichment analysis (GSEA) to assess the enrichment of its predicted targets. Finally, we kept TFs with adjusted p-values <0.05 and further identified MRs based on the ranking of the normalized enrichment score (NES). Our analysis found that a small series of TFs such as *Tcf4*, *Foxp4*, and *Mndal* emerged as top-MRs when core-MaSCs progress to luminal-MaSCs, whereas *Zeb2, Id3, Twist2*, and *Prrx1* are MRs for the transition from core-MaSCs to basal-MaSCs (Fig.5A and 5B). Interestingly, while some of these TFs are known to be expressed in MaSCs, others represent novel MRs in MaSC fate specification. Of particular interest are *Tcf4* and *Id3*, which have been implicated in development in other systems but not in MaSCs.

Straight *Tcf4* knockout animals are not viable^46^, consequently, whether they present mammary gland abnormalities is unknown. Homozygous *Id3*-KO mice are viable, and mammary abnormalities have been reported in mice lacking expression of other Id proteins. *Id4*-null mice exhibited reduced ductal and alveolar expansion^47^ and *Id2*-null mice showed impaired alveolar differentiation during pregnancy^48^. Individual *Id1* and *Id3* null strains do not present any phenotype due to compensatory mechanisms^49^. Unfortunately, *Id1* and *Id3* double knockout mice are embryonic lethal^50^ preventing further studies. Because these data illustrate the importance of ID proteins in the mammary gland we examined the expression of *Id* genes in our single-cell data (Fig. 5C). Remarkably, we found that although *Id4* expression is very high in mature basal cells, the expression of *Id3* is almost exclusive to basal-MaSCs. Furthermore, *Tcf4* is only expressed in non-basal-MaSCs.

**Fig. 5.**
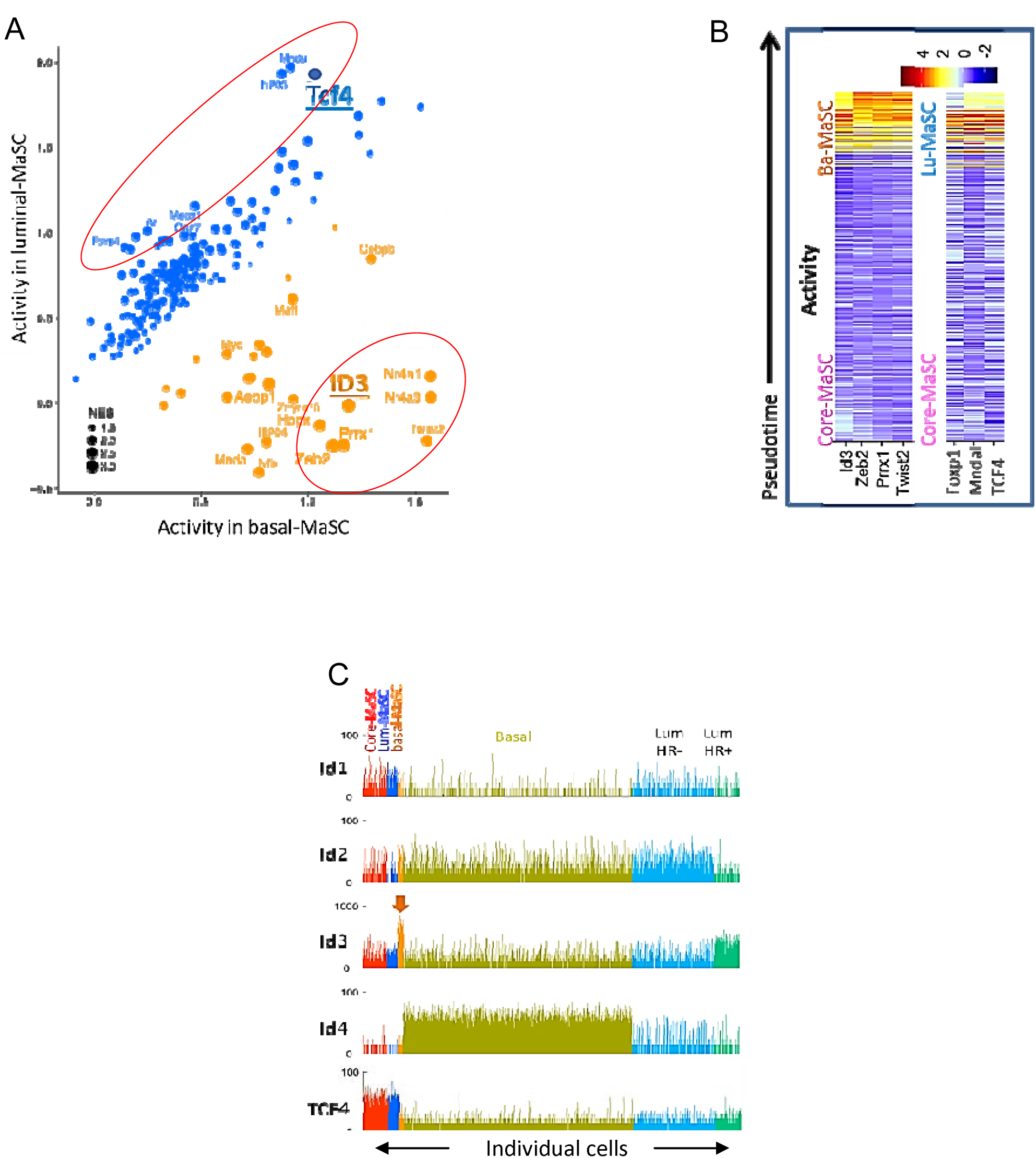
Master Regulators of MaSC subpopulations: A) NetBID activity analysis for TFs specifically active in basal-MaSC vs luminal-MaSCs. The dot size is proportional to the normalized enrichment score (NES) of the TF targets. B) Pseudotime analysis showing the change in the activity of MRs during the progression from core-MaSCs to basal-MaSCs or luminal-MaScs. C) Bar plot showing the expression of specific genes in mammary subpopulations (color-coded).

## DISCUSSION

The mammary epithelium undergoes continuous expansion and regression during the female lifetime^1–4^. These cycles of expansion need to be tightly regulated in time and space to ensure proper organ function and avoid pathological consequences. Although the existence of MaSCs supporting mammary epithelial expansion is well documented^5–9^, the mechanisms regulating homeostasis and differentiation of adult MaSCs are poorly understood. Characterizing these processes is critical for understanding the dynamics of the mammary epithelium and preventing and treating diseases such as breast cancer. However, conflicting results exist regarding the differentiation potential of MaSCs and while some studies have reported the existence of luminal and basal unipotent MaSCs^14^, others have found bipotent MaSCs that replenish both lineages^5,11–13^.

While previous studies have described the expression of protein C receptor (PROCR) as a marker of bipotent MaSCs^13^, our study goes one step further. Through the use of unbiased single-cell RNA sequencing (scRNA-seq) we have found that the PROCR+ MaSC population is composed of a core-MaSC subpopulation located at the top of the mammary epithelial hierarchy as well as two more lineage-committed subpopulations presenting basal and luminal molecular characteristics. Importantly, we have also delineated the activation pattern of the TFs during the early phases of lineage specification. Our studies indicate that *Tcf4* and *Id3* are two master regulators involved in the early acquisition of lineage-specific molecular features in MaSCs. The protein product of *Tcf4* called E2-2, belongs to the bHLH family of transcription factors that form homo- and heterodimers^51,52^. ID3 is also an HLH member; however, it works as an inhibitor of other bHLH proteins (including E2-2) by blocking their transcriptional activity^51,52^. Based on their anti-correlative activity observed in our preliminary data, we propose a model in which the acquisition of luminal features may be triggered by the expression of genes controlled by E2-2 whereas inhibition of E2-2 by ID3 leads to basal differentiation. Additionally, while ID3 and E2-2 are well-known partners, it is also possible that both of them work independently of each other. Thus, inhibition of other bHLH partners by ID3 and interaction of E2-2 with additional transcriptional regulators should also be considered. Of course, these alternatives are not mutually exclusive, and most likely, MaSC fate is influenced by a network coordinated by these two bHLH proteins consisting of common and specific downstream gene targets (Fig. 6). Thus, while our studies have unveiled *Tcf4* and *Id3* as some of the initial TF controlling lineage-specification, additional studies are needed to address the extension of this network and the importance of each hub.

**Fig. 6.**
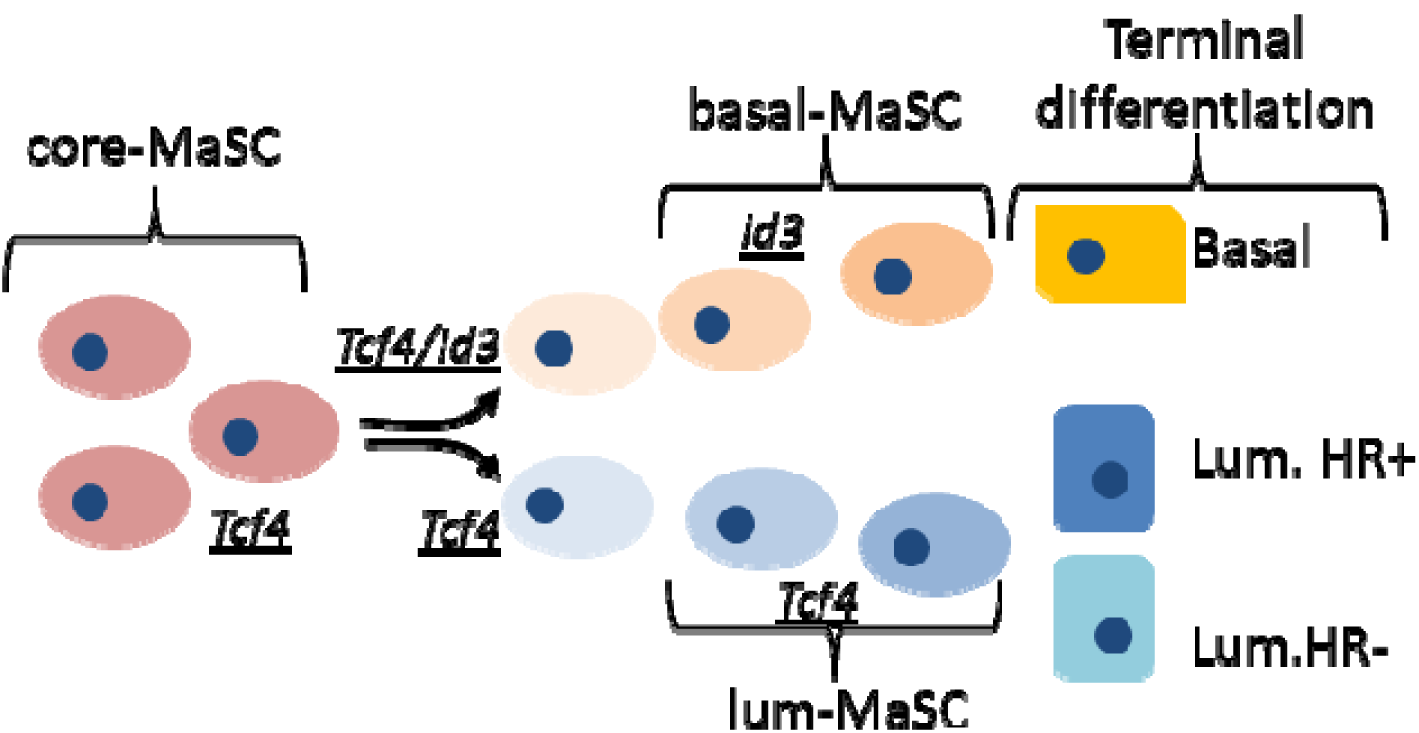
Proposed E2-2(Tcf4)/ID3 switch model: Bipotent adult core-MaSCs express high levels of the E2-2(Tcf4) transcription factor. During the early stages of lineage commitment, increasing expression of the HLH partner ID3 in some of the core-MaSCs inhibits Tcf4, and potentially other partners, leading to the acquisition of basal characteristics. In contrast, core-MaSCs that enter the differentiation process and do not express ID3 will acquire luminal lineage characteristics.

## Individual author contributions

JS and JY designed the research plan. KKY, EN, JS, and JY wrote the manuscript. EN, RW, DA coordinated the animal experiments and performed the mammary gland dissection and purification of mammary epithelial cells by FACS. KKY performed the computational and systems biology analysis. XD and LD developed the data portal. QP and CQ assisted single-cell RNA-seq data analysis. PM and AG provided additional support for animal care and mammary gland dissection. BJ, CR, BS, SN, and JE performed single-cell and bulk genomics profiling. YW, HS, and HC coordinated animal processing for single-cell studies.

## MATERIALS AND METHODS

### Mammary gland dissociation and epithelial enrichment (mouse)

#### Experimental animals

Wild-type BALB/cJ virgin female mice were obtained from The Jackson Laboratory. The mice were sacrificed at the specified age using a CO_2_ chamber. Animal maintenance and all experiments were performed under the Institutional Animal Care and Use Committee (IACUC) guidelines.

#### Determination of estrous cycle stage

Vaginal smears were taken by flushing with 15 μl of sterile PBS which were then spread on a slide and allowed to dry. The cells were visualized after staining with crystal violet and washing twice with deionized water. The stages were then identified as described (Byers *et al*., 2012).

#### Mammary gland dissociation and epithelial enrichment

The 3rd and 4th mammary glands were dissected from female mice, pooled, and minced followed by enzymatic digestion in DMEM/F12 media containing 2 mg/ml Collagenase A (Roche #11088793001) and 100 U/ml Hyaluronidase (Sigma #H3506) for 2 hours at 37°C. Single-cell suspensions were obtained via sequential incubations in pre-warmed 0.25% Trypsin-EDTA followed by 5 mg/ml Dispase II (Life Technologies #17105-041) in PBS with 0.1 mg/ml DNase I (Stemcell #07900), pipetting up and down at each step. Finally, the cells were incubated in Red Blood Cell Lysing Buffer (Sigma #R7757) and filtered through a 40 µm cell strainer. The samples were enriched for Lin- epithelial cells using the EasySep Mouse Epithelial Cell Enrichment Kit II (Stemcell #19758) according to the protocol.

#### Fluorescence-activated cell sorting (FACS)

The epithelial cell-enriched cell suspensions were incubated with the following antibodies: EpCAM-PerCP/Cy5.5, CD49f-APC, PROCR-PE, and Sca-1-BV421 (BioLegend #118220, #313616, #141504, and BD #562729). The LIVE/DEAD™ Fixable Aqua Dead Cell Stain (ThermoFisher #L34957) was used to determine cell viability. Antibody incubations were performed for 15 minutes on ice. The cells were sorted using a FACSAria II (BD) cell sorter using the following method: following size and doublet discrimination and viability screening, EpCAM+ cells were sorted for the Total Epithelium samples. To enrich for MaSCs, EpCAM+CD49fhighPROCR+ cells were sorted.

### Mammary gland dissociation and epithelial enrichment (Human)

Fresh tissue was obtained from mastectomy surgeries via the Biorepository Core at Mount Sinai according to an approved protocol. Samples were dissociated according to a published method (Shehata and Stingl, 2017). Briefly, the tissue was minced and enzymatically digested in DMEM/F12 media containing 2 mg/ml Collagenase A (Roche #11088793001), 100 U/ml Hyaluronidase (Sigma #H3506), 2% BSA, 5 µg/ml insulin, 50 μg/ml gentamycin, and 10mM HEPES overnight at 37°C. The samples were enriched for epithelial cells via centrifugation at 200xg for 5 minutes. Single-cell suspensions were obtained via sequential incubations in pre-warmed 0.25% Trypsin-EDTA followed by 5 mg/ml Dispase II (Life Technologies #17105-041) in PBS with 0.1 mg/ml DNase I (Stemcell #07900). The dissociated cells were subjected to FACS using the same method as above with the following antibodies: EpCAM-PerCP/Cy5.5, CD49f-APC, and PROCR-PE (BioLegend #324213, #313615, and #351903, respectively).

### Bulk RNA library preparation and sequencing

For samples with limited amount of RNA, we used NEBnext Single Cell/Low RNA Library prep Kit for Illumina to generate sequencing libraries (New England BioLabs). Libraries were sequenced on Illumina NextSeq 500 system (Illumina).

For samples with abundant RNA, stranded RNA libraries were prepared with 100ng total RNA using the KAPA RNA HyperPrep Kit with RiboErase (HMR) (Roche). RNA sequencing was performed on Illumina NovaSeq 6000 system (Illumina).

### Single Cell RNA Sequencing (scRNA-Seq) sample processing

Flow-sorted single cells were captured with the 10×Genomics Chromium Controller (10×Genomics). Libraries were prepared with the Chromium Single Cell Gene Expression 3’ v2 or v3 kits, and sequenced on either Illumina Hiseq 2500 or NextSeq 500 systems (Illumina). Sequencing data were run on Cell Ranger to generate feature barcode matrices and to perform clustering and gene expression analysis.

### Single cell RNASeq data analysis

Sequencing data were processed and converted to counts by Cell Ranger, the software tool developed by 10X genomics. Counts data were then converted to gene-by-cell expression matrix by Seurat ^1^. In brief, cells were filtered based on the unique number of genes and UMIs. Cells with UMI number and the unique number of genes too high or too low were filtered. Cells with a high fraction of mitochondrial genes were further filtered. The number of counts in the remaining cells were normalized by a scaling factor 100000 and gene expression level was quantified as E = log2(C+ 1), where C is the normalized counts for gene i in cell j. The resultant expression matrix was used for clustering analysis by our in-house pipeline scMINER. To take into account of the non-linear similarity between cells, scMINER quantified the distance between cells using mutual information. Leading MDS components of the distance matrix were used for identifying clusters using consensus clustering. Clusters were then manually curated using marker genes from literature. Comprehensive identification of marker genes in each cluster was done by differential expression analysis using Seurat.

### Developmental hierarchy and network inference

The developmental stages and pseudotime were inferred based on expression of individual genes using diffusion maps^2^ and the tool Monocle 2^3^. The ordering of cells based on diffusion maps and pseudotime was supported by the entropy measure, a metric to quantify the developmental plasticity, defined as -, where is the relative expression^4^. To better differentiate different populations, activities of various gene sets or signatures were used on top of marker genes expression. Activities of specific gene sets were calculated using NetBID^19,39^, which is essentially a weighted measure of expression of genes in the set. Network inference was performed using SJARACNE^18^.

In short, cluster-specific network was inferred based on the expression profile of the cells in the cluster based on mutual information dependency. Transcription factors with a high number of targets in the network were regarded as potential master regulators (MR). Gene set enrichment analysis (GSEA) and curated gene sets from MSigDB{Liberzon, 2011 #897 were used to identify potential biological functions of the targets. The activity of the potential regulators is calculated from the expression of its targets, and differential activities were inferred using NetBID. The regulators whose activities are correlated with the pseudotime were operationally defined as MR for the development.

## Notes

### Competing Interest Statement

The authors have declared no competing interest.

